# The rosy future paradox: Positive future thinking without task relevance enhances negative biases and anxiety for aversive events

**DOI:** 10.1101/2022.01.03.474768

**Authors:** Nicole D. Montijn, Lotte Gerritsen, Dana van Son, Iris. M. Engelhard

**Affiliations:** Department of Clinical Psychology, Utrecht University, Utrecht, The Netherlands; Institute of Psychology, Leiden University, Leiden, The Netherlands

## Abstract

Expectations have an important role in guiding behavior and the interpretation of novel information, but can contain negative biases such as is the case in anxiety disorders. Positive future thinking may provide an accessible way to attenuate these negatively biases. However, much is still unclear about the optimal form of such positive interventions, and it is unknown if the effects go beyond subjective experience. Here, we used a positive future thinking intervention to adapt the way a stressful event is experienced. Participants imagined either task-relevant (N = 21) or irrelevant (N = 21) positive future events before being subjected to the Trier Social Stress Task, or did not receive the intervention (N= 20). We recorded resting state EEG during the anticipation and recovery phases of the TSST to assess intervention and trait anxiety related differences in the level of frontal delta-beta coupling, which is considered a neurobiological substrate of emotion regulation. Results show that the intervention reduces event-related stress and anxiety, and increases social fixation behavior and task performance, but only if future thinking is task relevant. Paradoxically, task-irrelevant positive future thoughts enhance negative perceptual biases and stress reactivity. This increase in stress reactivity in the task-irrelevant group was corroborated by the elevated levels frontal delta-beta coupling during event anticipation, especially for those with high trait anxiety. This suggests an increased demand for emotion regulation following the task-irrelevant intervention, possibly due to the contextual incongruence between positive imagery and the stressor. Together, these results show that positive future thinking can mitigate the negative emotional, behavioral and neurobiological consequences of a stressful event, but that it should not be applied indiscriminately. Task-relevant positive future thinking can be an accessible way to boost efficacy of exposure therapy for pathological anxiety, and can help people deal with negative anticipation in daily life.

## INTRODUCTION

Anticipatory anxiety and stress are as much a part of everyday life as they are of certain mental disorders. Such anxiety is associated with negatively biased expectations and interpretations that can cause emotional distress (Butler & Mathews, 1987; Mathews & MacLeod, 2002). Expectations have an important role in guiding behavior and the interpretation of novel information. They are typically shaped by prior experience (Gilboa & Marlatte, 2017), which provides a perceptual filter that influences attention (Hutchinson & Turk-Browne, 2012; Ryan & Shen, 2020), perception (Dijkstra et al., 2021; Mather & Sutherland, 2011) and ultimately memory (Audrain & McAndrews, 2020; Masís-Obando et al., 2021). For example, you may avoid a shorter route to work because it is notoriously high in traffic, or perceive a compliment from a typically unfriendly co-worker as insincere. As such, negative expectation does not just protect an individual from threat but also helps moderate emotional responses to unpleasant situations, or prevent them altogether (Miloyan et al., 2016). However, it can bias processing of novel experiences, so negativity can over time become disproportionate to the situation (Clark & Beck, 2010). For example, in the case of the co-worker, their attempts at making a genuine connection may go unnoticed, preventing potential positive interactions.

Expectations of an event may be expressed and evaluated through episodic future thinking. This involves the mental simulation, or pre-experiencing, of future events by recombining elements from previous experiences (Addis et al., 2008). Future thinking is important for a range of cognitive functions including planning, likelihood estimation, decision making, and emotion regulation (Schacter et al., 2017). Its role in emotion regulation is also reflected on a neural level. Future simulation relies on functional connectivity between the hippocampus and prefrontal cortex during event construction (Benoit et al., 2014; Campbell et al., 2018; Demblon et al., 2015), an area that has been related to emotion regulation processes and cognitive control. Because episodic future thinking takes a role in both the expression of anticipatory bias and the regulation of the accompanying emotional response, it may provide an accessible way to attenuate negatively biased cognitions.

Indeed, recent work shows that future thinking can positively bias the interpretation of neutral narratives (Devitt & Schacter, 2018), and inhibit the recollection of contextually similar scenarios (Ditta & Storm, 2016). Furthermore, increasing the level of episodic detail with which future events are simulated enhances emotion regulation strategies and improve psychological well-being towards worrisome future events (Jing et al., 2016, 2017). Beyond future thinking, positive imagery interventions have been developed to reduce anticipatory anxiety and stress (Pile et al., 2021). Interventions that use personally or contextually relevant imagery appear to produce consistent effects (e.g. Landkroon et al., 2021; Renner et al., 2019). However, it is unclear whether they work ubiquitously, independent of context or trait predisposition, and if these effects go beyond subjective experience. Of particular interest is whether task-relevance is indeed a boundary condition for the effect of positive interventions, and whether trait anxiety may limit efficacy as it is associated with deficits in emotion regulation (Cho et al., 2019; Liu et al., 2018).

Here, we addressed those questions by having participants imagine positive future events before being subjected to the Trier Social Stress Test (TSST; Kirschbaum et al., 1993), which is an aversive task that involves an impromptu presentation in front of a jury panel. To address the notion that efficacy might depend on task-relevance of the intervention, we compared no-intervention controls to participants who imagined either positive task-relevant or task-irrelevant future events. The goal was to mitigate the negative emotional response that is generally triggered by the TSST using positive future thinking, and skew subjective perception towards more positive interpretations of this stressful task. We expected both intervention groups to benefit from the intervention compared to controls, but expected that the task-relevant group would show the most improvement. We used a combination of self-report measures, eye tracking, and electroencephalography (EEG) to assess intervention-related differences in stress reactivity and emotion regulation.

Emotion regulation depends on connectivity between the pre-frontal cortex and limbic areas, like the hippocampus and amygdala (Banks et al., 2007). Cross-frequency coupling, or the interaction between two different neural oscillation frequencies, can be used as a measure for such functional connectivity (Canolty & Knight, 2010). Of interest is the level of coupling between delta (1 – 4 Hz), associated with affective processing and anxiety (Knyazev, 2007; Knyazev et al., 2005), and beta (14 – 30 Hz) oscillations, associated with cognitive control (Buschman & Miller, 2007; Engel & Fries, 2010). Specifically, frontal delta-beta phase-amplitude coupling (PAC) has been proposed as a neural marker for emotion and stress regulation (Schutter & Knyazev, 2012; Schutter et al., 2006). Earlier reports, that used a similar task as the current study, showed that delta-beta PAC typically increases when state nervousness and anxiety increase (Knyazev, 2011; Poppelaars et al., 2018). Therefore, delta-beta PAC could be a viable measure for differences in stress and emotion regulation.

## RESULTS

We randomly allocated 62 participants, none of whom self-reported current psychiatric impairment, to one of three intervention groups; the task-irrelevant, task-relevant and a no-intervention control group. The **task-irrelevant** group was asked to imagine 15 generally positive episodic future events (e.g., going on vacation), while the **task-relevant** group imagined 15 positive episodic future events in which they were giving a presentation (e.g., a successful thesis defense) before being subjected to the Trier Social Stress Test (TSST). During the TSST, participants had to give an impromptu 5-minute presentation about climate change in front of a jury panel, and were told that they would be rated on their performance. We assessed intervention related changes in stress and anxiety using a combination of behavioral measures and resting state EEG. EEG data are described for three phases of the experiment: pre-intervention **baseline**, pre-presentation **anticipation** and post-presentation **recovery**. Additionally, we assessed whether the intervention enhanced performance and task engagement (eye-tracking) during the TSST presentation.

### Subjective measures

#### No baseline group differences

Participants started with a baseline assessment of state and trait anxiety levels (STAI; Spielberg et al., 1970), as well as trait based levels of worry (PSWQ; Meyer et al., 1990), stress reactivity (PSS; Cohen et al., 1983) and imagery ability (VVIQ; McKelvie, 1995). Group comparisons (see Table 1) revealed no baseline differences on any of the measures: trait anxiety (STAI-T), *F*(2, 61)= .073, *p* = .93, state anxiety (STAI-S), *F*(2, 61)= 1.07, *p* = .350, worry (PSWQ), *F*(2, 61)= .092, *p* = .912, stress reactivity (PSS), *F*(2, 61)= 1.939, *p* = .153, and visual imagery ability (VVIQ), *F*(2, 61)= 1.000, *p* = .374. This suggests that the randomization of participants across conditions was successful. Following the baseline measures, participants completed either the task-relevant or task-irrelevant positive future thinking intervention, or, in case of the control group, proceeded directly to the TSST.

**Table 1.**
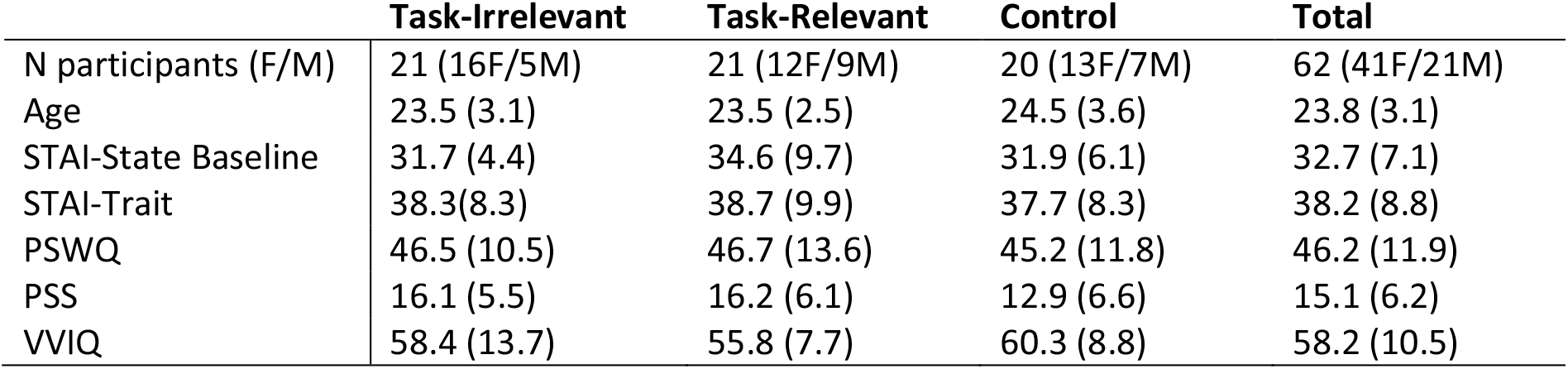
Demographics and Baseline Questionnaire scores (Means and SD) per experimental group.

#### Task-irrelevant positive future thinking leads to more stress reactivity during anticipation

After the positive future thinking intervention, participants were instructed to prepare a 5-minute presentation on climate change that they would have to present in front of a jury panel. They were given 2 minutes to study a fact sheet on this topic, and another 5 minutes of mental preparation time (see Figure 1B). After this anticipation phase, we assessed subjective levels of anticipatory stress using the Primary Appraisal (situational threat and challenge) and Secondary Appraisal (situational control and self-confidence) scale (PASA). A positive Stress Index (primary – secondary) reflects that the perceived threat or challenge outweighs the perceived ability to control the situation. We expected that both intervention groups would have a lower Stress Index than the no-intervention control group, and that anticipatory stress would be lowest in the task-relevant group. However, we found a significant main effect of group, *F*(2, 59)= 4.815, *p* = .012, *η*_*p*_^*2*^ *=* .14, that was driven by a higher Stress Index in the task-irrelevant group compared to both the task-relevant and control group (see Figure 1C). This suggests that task-irrelevant positive future thinking, before being exposed to an aversive task, increases anticipatory stress instead of helping to reduce it.

**Figure 1.**
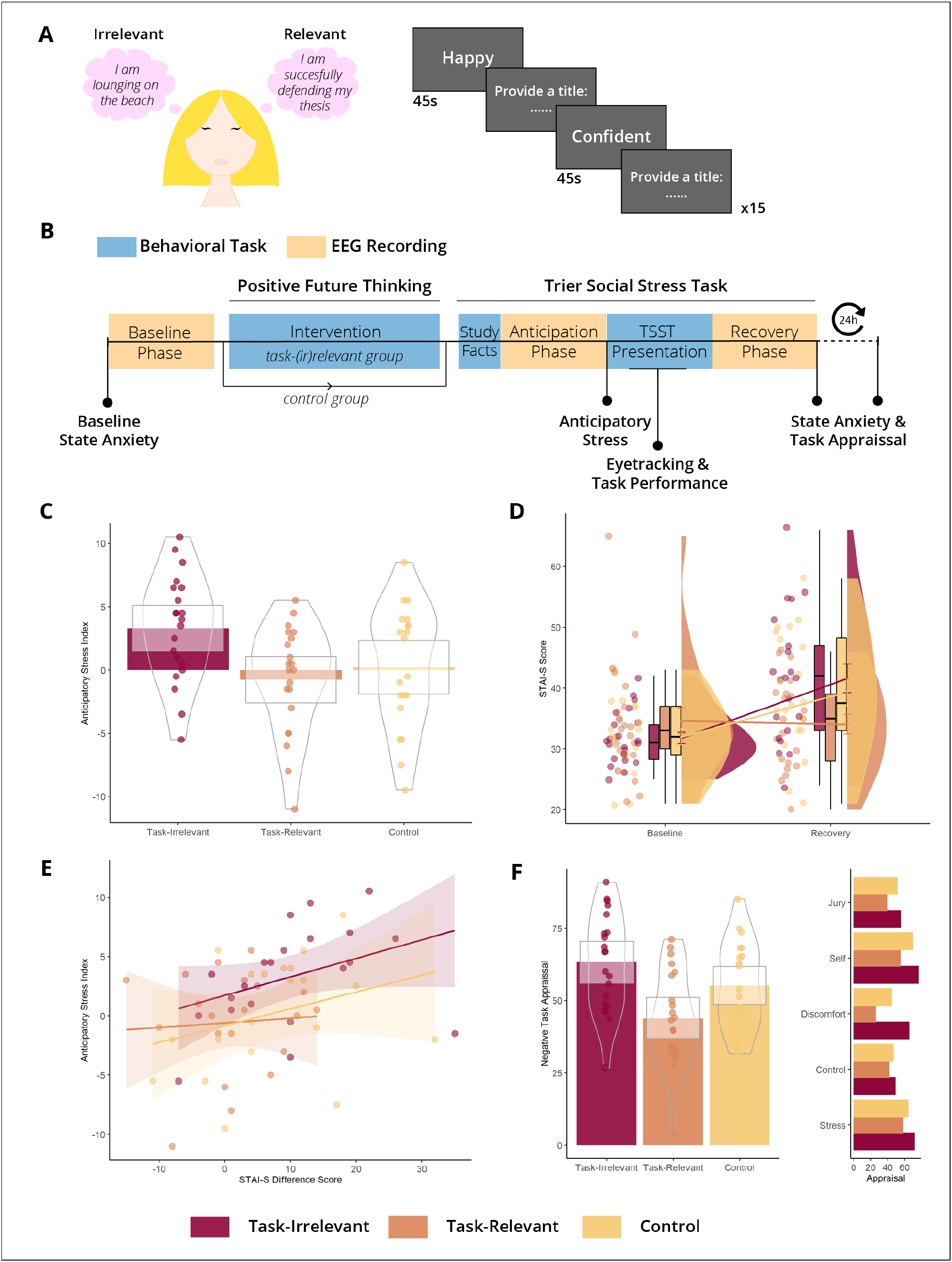
Design and Subjective Data. **A)** Task design Positive Future Thinking intervention. Participants either imagined 15 generally positive future events (Task-Irrelevant Group) or 15 positive future events where they gave a presentation (Task-Relevant Group). Positively valanced cue words were presented to aid event construction. Participants were asked to provide a title for every imagined event to ensure task-compliance. **B)** Study design including timepoints at which specific measures were taken. **C)** Response distribution for Anticipatory Stress Index. Colored bars represent group mean, white squares show 95% CI, violins and points show distribution of individual participants. Positive scores reflect maladaptive stress while negative scores reflect adaptive stress. **D)** Between groups response distribution and group means of STAI-S scores at Baseline and post-TSST Recovery. Graph shows an increase in State Anxiety for the Task-irrelevant and Control group, but not for the Task-relevant group. Scores on the STAI-S range from 20 to 80. **E)** Correlation between Anticipatory Stress levels and task-related changes in State Anxiety. **F)** Subjective task appraisal scores separated by experimental group. Bars represent mean scores, violin and points show response distribution, boxes represent 95% CI. To the right, appraisal scores are separated by both experimental group and appraisal sub-scale. Scores ranged from 0 (not negative at all) to 100 (very negative).

#### Task-relevant positive future thinking prevents event-related negativity and anxiety

Directly after the TSST, participants completed the STAI-S again to assess changes in State Anxiety following their presentation. A 2 (time: Baseline, Recovery) x 3 (Group: Task-irrelevant, Task-relevant, Control) mixed analysis of variance (ANOVA) revealed a significant interaction between time and group (see Figure 1D), *F*(2, 59)= 5.914, *p* = .005, *η*_*p*_^*2*^ *=* .167. This interaction was explained by an increase in State Anxiety in the Control, *t*(19) = -3.263, *p* = .004, *d* = .72, and Task-Irrelevant group, *t*(20) = -4.462, *p* = .000, *d* = .93, but not in the Task-Relevant group, *t*(20) = .243, *p* = .81, *d* = .05.

Participants also completed an 11-item questionnaire that assessed their subjective appraisal of their performance and the event itself (i.e. the TSST presentation). Items were rated using a slider scale ranging from 0 to 100, with higher scores reflecting more negative appraisal. Items were divided into five categories: Experienced Stress, Situational Control, Discomfort, Self Evaluation and Jury Evaluation. Experimental groups differed significantly from each other in overall task appraisal (see Figure 1F), *F*(2, 59) = 7.608, *p* = .001, *η*_*p*_^*2*^ *=* .205. Pairwise comparisons (Bonferroni corrected) showed that the Task-Relevant group (*M* = 43.9, *SD* = 3.5) appraised the task significantly more positively (*p* = .001) than the Task-Irrelevant group (*M* = 63.3, *SD* = 3.5), and at a trend level (*p* = .088) compared to the Control group (*M* = 55.2, *SD* = 3.6). Analysis of the five sub-categories showed that the Task-relevant group scored consistently lower (i.e. more positive appraisal) on all categories (see Figure 1F), but significant group differences were only found for the Discomfort, Self Evaluation and Jury Evaluation sub-scales (all *p* < .01). These effects persisted in the follow-up measurement 24 hours later.

### Behavioral Measures

#### Task-relevant positive future thinking boosts task performance

To assess whether the intervention enhanced task performance, we calculated the number of climate change facts that participants remembered to incorporate in their presentation. We found a significant effect of group *F*(2, 59) = 3.481, *p* = .037, *η*_*p*_^*2*^ *=* .11, that was explained by the task-relevant group (*M* = 6.67, *SD* = 1.35) presenting significantly more facts than the control group (*M* = 5.15, *SD* = 2.01), *t*(39) = 2.849, *p* = .007, *d* = .89. However, contrary to what might be expected based on the subjective data, the task-relevant group did not present more facts than the task-irrelevant group (*M* = 6.1, *SD* = 2.11), *t*(39) = -1.041, *p* = .304, *d* = .46.

#### Task-relevant positive future thinking increases social fixation behavior

Finally, we examined whether the intervention positively impacted fixation behavior during the TSST presentation. Avoidance of eye-contact is seen as a behavioral correlate of social anxiety (Chen et al., 2020; Herten et al., 2017). Therefore, we expected participants who experienced more stress to fixate less on faces and more on the surrounding environment. In line with the state anxiety, task appraisal, and task performance measures, the Task-Relevant group fixated significantly more on faces of the jury members than the Task-Irrelevant group, *t*(35) = 2.097, *p* = .043, *d* = .69, and tended to fixate more on faces than the Control group, *t*(34) = 1.828, *p* = .076, *d* = .61 (see Figure 2). This suggests that the Task-Relevant group experienced less social stress than both the Task-Irrelevant and Control group, as gaze avoidance is seen as a common strategy to reduce social stress or discomfort.

**Figure 2.**
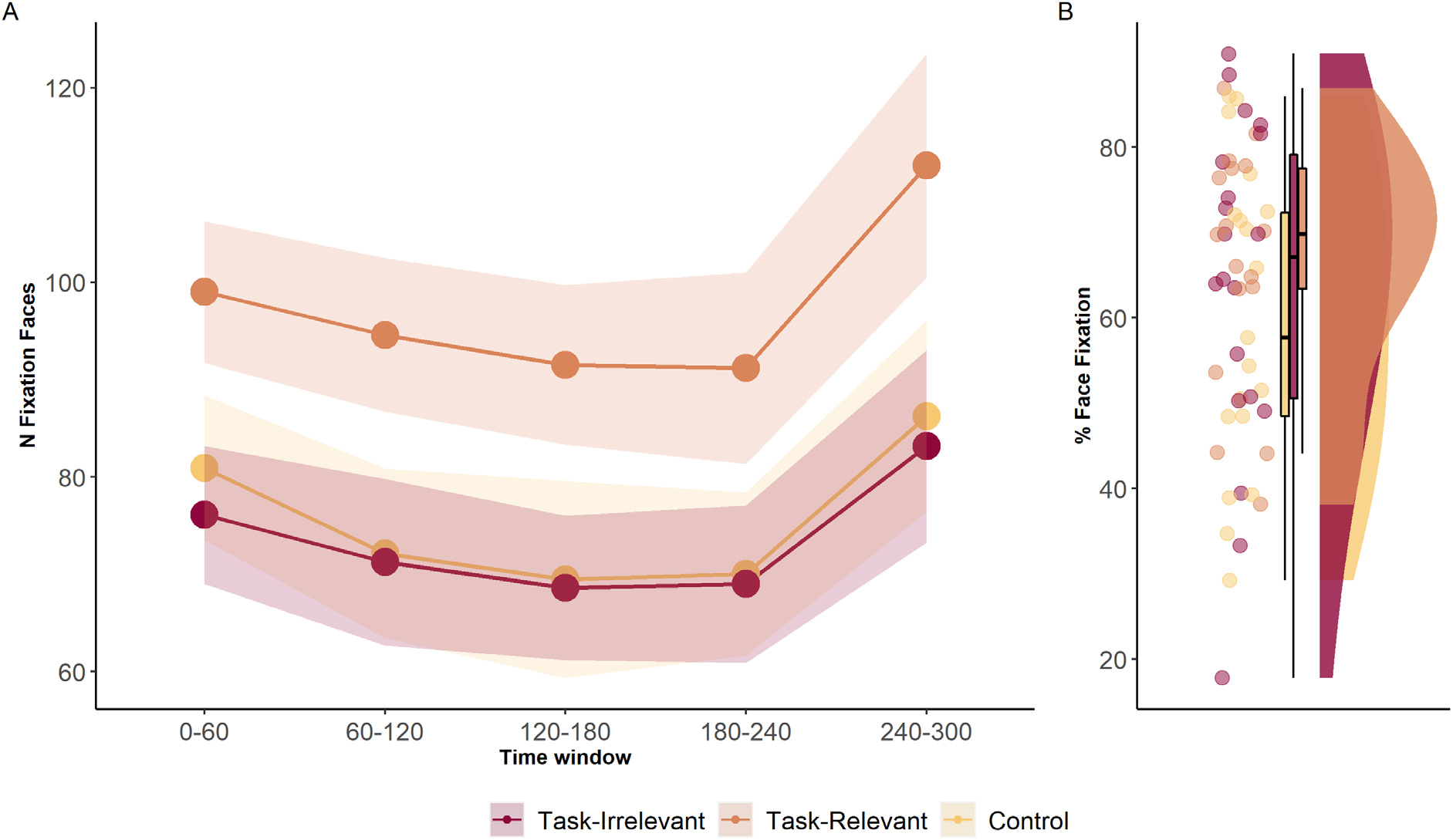
Fixation data during TSST. **A)** Mean (shaded areas show SE) number of fixations on faces of the three jury members during the TSST presentation divided by experimental group and 60sec time window. **B)** Percentage of fixations on faces out of the total number of fixations per participant.

### Cross-frequency EEG Measures

#### Delta-beta dPAC increases as a function of trait anxiety and stress

The assumed mechanism behind the positive future thinking interventions is that they promote emotion regulation ahead of the stressful event. Therefore, we expected that levels of frontal delta-beta phase-amplitude coupling (see Figure 3A), a measure for stress and emotion regulation, would increase more for the intervention groups (Task-Relevant > Task-Irrelevant) leading up to the TSST compared to the control group, and wind down again during recovery. To assess this, we ran two linear mixed regression models, one for the upward slope (early baseline to late anticipation) and one for the downward slope (late anticipation to late recovery), using group and time as fixed factors and allowing random slopes per subject. Compared to the control group, delta-beta dPAC levels in the task-irrelevant group increased from early baseline into late anticipation (β = .11, *p* = .09) and decreased again from late anticipation to late recovery (β = -.17, *p* = .07) (see Figure 4A). This effect was not found for the Task-Relevant group.

**Figure 3.**
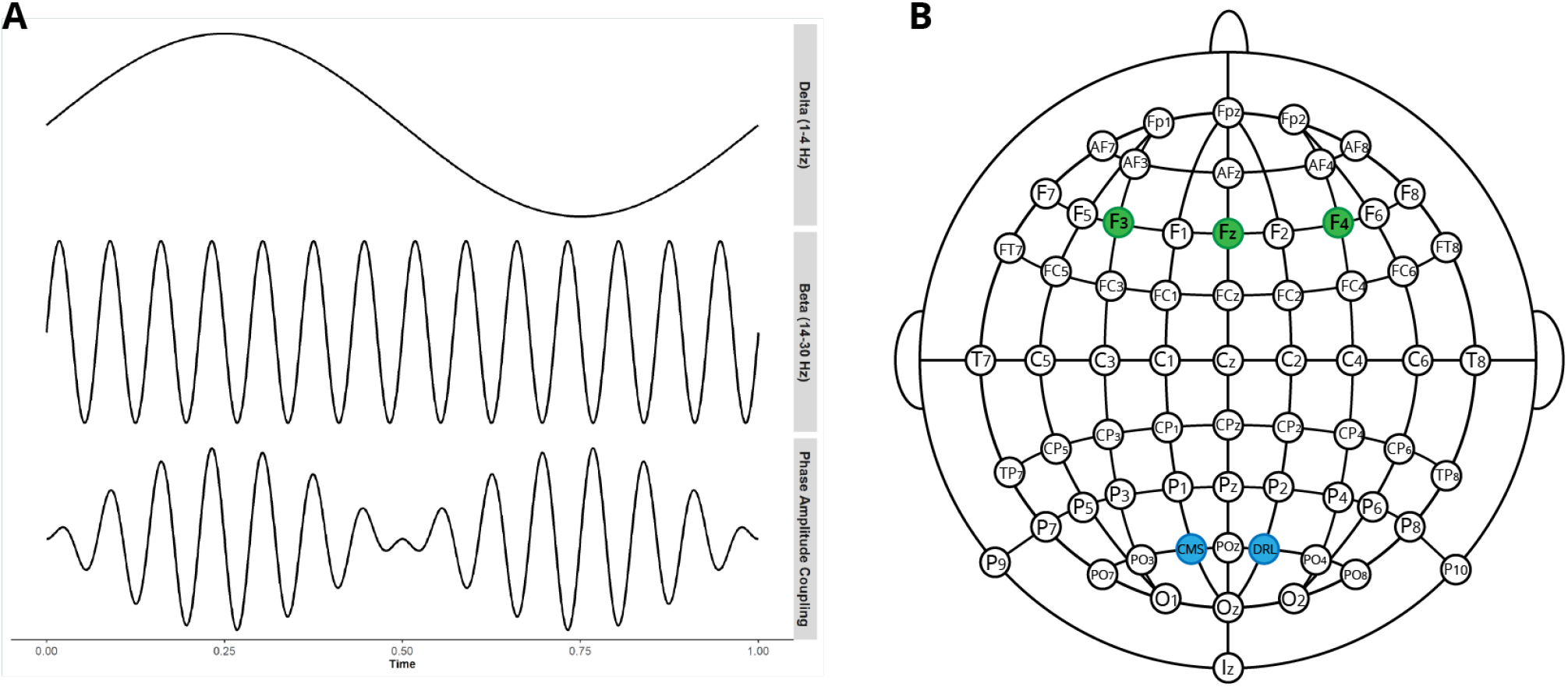
Cross frequency coupling. **A)** Illustration of how the phase of low-frequency delta oscillations modulate the amplitude of high-frequency beta oscillations, or delta-beta phase-amplitude coupling. **B)** Lay-out of 64-electrode Biosemi EEG cap. The analyses used the averaged z-scored signal of F3, Fz and F4 (in green). Ground (DRL) and online reference (CMS) electrodes are depicted in blue. Data were re-referenced to the average of all 64 electrodes.

**Figure 4.**
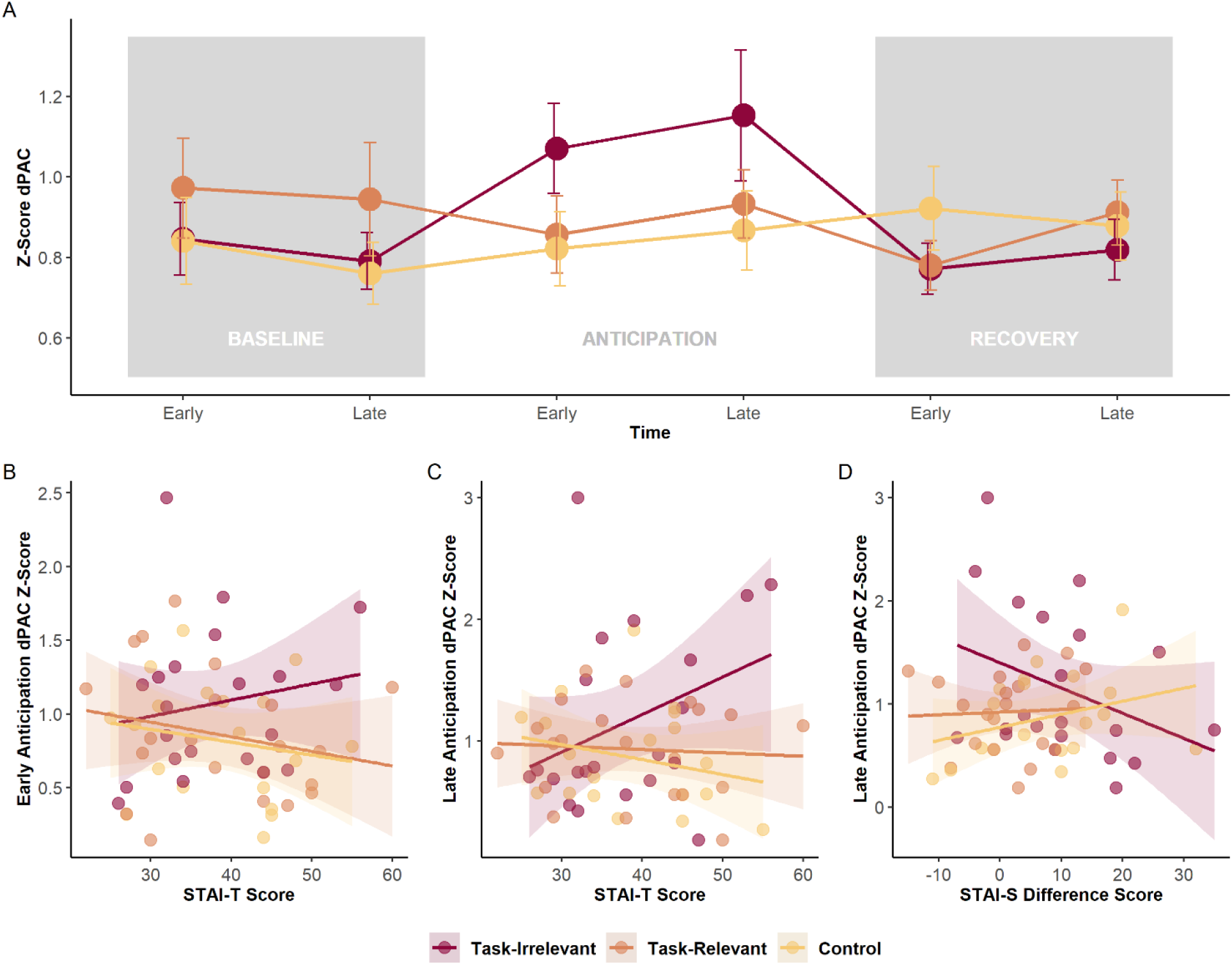
Delta-beta phase amplitude coupling. **A)** Mean and SE of Z-scored delta-beta dPAC (average of electrodes Fz, F3 and F4) per experimental group. Line graph shows the progression of dPAC levels over the three phases of the experiment (Baseline, Anticipation and Recovery). Early and Late reflect an average dPAC level for the first (early) and last (late) 80 seconds of that phase. **B)** Scatter plot showing group differences in the correlation between Early Anticipation dPAC and Trait Anxiety scores. **C)** Correlation between Late Anticipation dPAC and Trait Anxiety scores **D)** Correlation between Late Anticipation dPAC and State Anxiety Difference scores (Recovery – Baseline).

Next, to assess the effect of trait vulnerability on emotion regulation and the interaction with the intervention groups, we included Trait Anxiety as a fixed effect in both models. Leading up to late anticipation, we found a significant interaction between group, time and trait anxiety levels, *F*(2, 178.94)= 3.053, *p* = .04 (see Figure 4B and C). Trait anxiety significantly predicted increases in dPAC over time in the Task-Irrelevant group (β = .019, *p* = .014). From late anticipation to late recovery, higher levels of trait anxiety were also associated with steeper decreases in dPAC in the Task-Irrelevant group (β = -.02, *p* = .07). Given that trait anxiety is generally associated with ineffective emotion regulation and the high levels of anticipatory stress in the Task-Irrelevant group, this prompts the question whether delta-beta PAC might code for stress reactivity rather than emotion regulation.

To test this, we used delta-beta dPAC scores during late anticipation as a predictor for changes in State Anxiety levels pre to post TSST (Recovery minus Baseline). Increased delta-beta coupling in the Task-Irrelevant group was negatively predictive of task-related increases in State Anxiety (β = -12.3, *p* = .06). This suggests that dPAC is indeed a marker for emotion regulation, as individuals with higher dPAC levels in anticipation of a stressor subsequently reported little to no increase in state anxiety levels (see Figure 4D). The effect was found despite generally higher state anxiety and stress in the Task-Irrelevant group (see Figure 1C and D).

## DISCUSSION

In this study, we investigated if positive future thinking can be used to attenuate negatively biased perception of a social stressor (TSST). Our data show that task relevance of the intervention, and not positivity alone, determines its benefit. The task-relevant group did not increase in state anxiety as a result of the TSST, while both the control and task-irrelevant group did. Furthermore, the task-relevant group showed more positive appraisal about the TSST, had better task performance, and engaged in more eye contact with the jury members compared to the control group. Positivity without task relevance, on the other hand, led to more negative task appraisal and a more severe stress reaction in response to the social stressor. This adverse effect of the task-irrelevant positive intervention may have been somewhat counteracted by the engagement of neural stress regulation mechanisms. Participants in de task-irrelevant group, in particular those with higher levels of trait anxiety, showed increasing levels of frontal delta-beta phase amplitude coupling in the period leading up to the stressor. These elevated levels of delta-beta PAC were negatively predictive of task-related increases in state anxiety. This suggests that individuals who engage this neural system are generally more successful at reducing the negative emotional consequences of enduring an aversive event.

Our hypotheses regarding the working mechanism of the interventions were centered on the notion that positive future thinking promotes effective emotion regulation, as a way of planning ahead (i.e. the event will still be stressful but participants feel more in control). Instead, we found that when the intervention is task-relevant, it prevents the anticipatory stress reaction that the TSST typically induces. Therefore, instead of *ad hoc* emotion regulation, task-relevant positive future thinking may facilitate the *a priori* re-appraisal of aversive events.

A possible explanation for the discrepancy between task-relevant and irrelevant future thinking in terms of stress response is that the intervention modulates memory schema activity. Higher-order conceptual knowledge is used as a perceptual filter that facilitates the interpretation of novel information as well as the selection of appropriate actions (Gilboa & Marlatte, 2017; Wang & Morris, 2010). Through future thinking, the task-relevant group may have instantiated contextually appropriate positive schema, which likely caused the interpretation of the new external information to assimilate into the active schema (Gilboa & Moscovitch, 2017; Tse et al., 2007). In the task-irrelevant group, the instantiated schema did not hold contextually relevant information that could help the individual deal with the stressor. Such incongruency may have prompted the reinstatement of the dominant schema for having to give a presentation in front of a jury (Frühholz et al., 2011; van Kesteren et al., 2020), which is negatively biased for many people. Furthermore, this incongruence resulted in that these participants felt less prepared to deal with the stressor, as reflected by heightened anticipatory stress and stronger engagement of frontal emotion regulation. Participants in the control group were also negatively affected by the stressor, but may have felt relatively more prepared during anticipation, because they were not distracted with irrelevant positive information beforehand.

This study has important implications and recommendations for psychological interventions that leverage positive imagery or simulation-based learning to reduce anxiety. First, positivity should not be applied indiscriminately. General positivity training may be beneficial in affective disorders like depression (Boland et al., 2018), where overall positivity tends to be reduced (Bjärehed et al., 2010), but it does not help to reduce anticipatory anxiety and (paradoxically) increases it. As data from our task-irrelevant group shows, by replacing negatively biased event anticipation with irrelevant positive information, the individual may feel less equipped to handle the situation. Second, the task-relevant positive future thinking intervention is effective even if it does not specifically address the upcoming stressful situation. That is, our task-relevant group imagined scenarios where they gave a presentation, but the presentations could concern anything from a wedding speech to a job application. This might be helpful in situations where individuals have a difficult time envisioning positive alternatives to a highly feared situation. Finally, due to its positive effects on anticipatory stress and task performance, task-relevant positive future thinking could be a useful tool to enhance both exposure willingness and efficacy in pathological anxiety (also see Landkroon et al., 2021).

This work extends findings on the transfer of valence through simulation-based learning. A recent study showed that future thinking can serve as a substitute for lived experience in updating pre-existing beliefs or attitudes. By imagining liked (or disliked) people together with a neutral location, participants changed their appraisal of the location towards the valence associated with the person (Paulus et al., 2021). Here, we show that such transfer of valence does not just apply to existing semantic representations, but also immediately affects the interpretation of new experiences that are semantically related. These positive effects on event appraisal remain at 24-hour follow-up. So, while it is unclear whether future thinking can establish long-term changes in pre-existing beliefs, the effects on information learned directly following the intervention are relatively stable.

Our findings also extend earlier work on the relation between stress and frontally mediated delta-beta coupling. Functionally, delta-beta PAC has been positioned as a stress regulation mechanism. This assumption is fueled by previous research (Knyazev, 2011; Poppelaars et al., 2018) showing that coupling increases as a function of state nervousness during anticipation (but see Poppelaars et al., 2021), which our results corroborate. However, these earlier studies do not include measures that reflect whether higher PAC is associated with more effective regulation of stress, merely that PAC increases as a response to stress. This study shows that people with higher levels of delta-beta PAC during anticipation have a lower increase in state anxiety following the stressor. So, while replication of these findings is needed, our data indicate that delta-beta PAC is indeed reflective of adaptive stress regulation. However, as noted by others (Brooker et al., 2015; Harrewijn et al., 2016; Poppelaars et al., 2021), whether this mechanism is actually engaged seems to depend both on context and trait predisposition.

Specifically, only participants in the task-irrelevant group experienced the levels of stress that prompted the engagement of this stress regulation mechanism (i.e. frontal delta-beta PAC). In addition, trait anxiety was positively correlated to elevations in delta-beta coupling during both anticipation and recovery. Similar moderating effects of trait anxiety have been found for other slow/fast wave EEG patterns, like delta-beta and theta-beta ratio, in relation to stress regulation and threat bias (Angelidis et al., 2018; Putman, 2011; van Son et al., 2018). These data should be interpreted with caution, but may suggest that some individuals with a trait vulnerability engage this frontal delta-beta PAC system to regulate stress when cognitive resources that help deal with the situation are low. It should be noted that increased activity does not automatically mean that emotion regulation is also effective (Kret et al., 2011; Sylvester et al., 2012), as our data also suggest. Highly anxious individuals may exert more effort to regulate their emotions in response to stress, but vary in their level of efficiency in doing so. This was also shown in an IAPS picture viewing task, in which individuals with high trait anxiety showed greater engagement of prefrontal emotion regulation systems to establish similar levels of emotional down-regulation as low-anxiety individuals (Campbell-Sills et al., 2011).

To summarize, positive future thinking can be an effective tool to induce *a priori* positive reappraisal of aversive situations and enhance task performance and goal-directed behavior. However, the contextual relevance of the imagined future scenarios to the aversive event is a clear boundary condition for this effect: when future thinking is incongruent with the aversive event, it can (paradoxically) increase stress and anxiety. Task-relevant positive future thinking may be used to increase willingness and efficacy of exposure therapy for pathological anxiety, and could be an accessible way for people to deal with negative anticipation in daily life. Over time, this could help to update the negative biases surrounding these situations, but this is an empirical question that awaits future research.

## METHODS

### Participants

We tested 65 students recruited at the Utrecht University campus (see Table 1 for demographics), none of whom reported any current psychiatric impairment. Furthermore, female participants were required to be on hormonal birth control to control for potential bias in the hormonal stress response due to fluctuations of female hormones (Espin et al., 2013). Participants were randomly assigned to one of three experimental groups: task-irrelevant positive, task-relevant positive and control. All participants provided written informed consent. They were remunerated with money or course credit for their participation. The study was approved by the institutional ethical review board at Utrecht University (FETC19-053).

Three participants were excluded from the entire analysis due to technical problems that forced us to quit the test session before all essential tasks had been performed. This led to a total sample size of 62 before data pre-processing (21 task-irrelevant, 21 task-relevant, and 20 control). Furthermore, 6 participants (4 task congruent, 1 task-irrelevant and 1 control) were excluded from the eye-tracking analyses due to large amounts of missing data (see Eye tracking pre-processing for further details on exclusion criteria), and 3 participants (one of each group) were excluded from the EEG analysis due to poor data quality.

### Procedure

Data acquisition took place over two consecutive days. The first session took place in the lab between 12:00 and 18:00 to limit variability in stress reactivity due to the circadian cortisol rhythm. The session took about 1,5 hours to complete. Participants started by filling out a battery of trait and state questionnaires (see Baseline Self-Report) followed by 5 minutes of eyes closed resting state EEG. This was followed by the positive future thinking intervention (see Positive Future Thinking), and then the Trier Social Stress Test (see TSST). For the control group, the order of these two tasks was reversed. Regardless of experimental group, the TSST was always directly followed by a questionnaire on task appraisal and memory for the preceding event (see Task Appraisal Self-Report). For the control group, an additional measure of this questionnaire was taken right after the future thinking intervention. However, for the purposes of this study the retroactive effect of positive future thinking on task appraisal (i.e. the additional task-appraisal measurement in the Control group) was not considered.

The second session consisted of a follow up questionnaire that consisted of the same items as the Task Appraisal Self-Report and a debriefing. This session was completed at home.

#### Positive Future Thinking

Participants were subjected to a positive future thinking intervention either before (task-irrelevant and task-relevant) or after (control) the TSST. Participants imagined fifteen positive future events that could occur within the next five years of their lives in response to a cue word. Participants were instructed to imagine the event in as much detail as they could, and to envision scenarios that evoked a highly positive emotion. Participants in the task-relevant and control groups received the additional instruction that the future events should involve them giving an oral presentation in front of a group of people (two or more). To allow more diversity between scenario’s any event involving some type of public speaking, such as receiving an award or a thesis defense, was accepted as ‘presenting’ as long as they were the one speaking. Participants in the

Task-Irrelevant group were simply instructed to imagine positive events that could occur in their personal future, such as going on vacation or a fun day with friends.

Cue words were selected from a subset 25 positively valanced cue words that are commonly used as part of the Autobiographical Memory Task (Williams & Broadbent, 1986). An independent student sample (N = 21) rated all 25 words on subjective valence and arousal, as well as ease of simulation for both generally positive events and events involving an oral presentation. Words that rated highest across all four measures were selected for use.

The task started with two practice trials to familiarize participants with the procedure. Practice trials were 3 minutes each. During that time participants were asked to imagine a detailed positive future event in response to the cue word that was presented on screen and describe the event to the experimenter as vividly as they could. If necessary, the experimenter would use general probes from the Autobiographical Interview to elicit a more specific or detailed account. For the remaining 13 trials, per trial one cue word was presented in the middle of the screen for 45 seconds, during this time participants had to construct a positive future event and elaborate on it as much as they could to create a vivid mental image. Next, participants were asked to type a short title (3 – 5 words) for the event to ensure task compliance. Trials were separated by a 5 second break, and a longer break of one minute halfway down the task. The task took about 25 minutes to complete.

#### TSST

The presentation part of the Trier Social Stress Test was used as the aversive episodic event that all participants were subjected to. Right before onset of the task, participants were informed that they would have to give an impromptu 5 minute presentation as if it were a job interview in front of a jury panel, whom they would be able to see through a video call. Participants were led to believe that the jury panel would be evaluating both their presentation and behavioral characteristics, and that their entire presentation would be recorded for subsequent analysis. In reality the ‘video call’ was a prerecorded video. To further standardize the presentation, all participants had to give a presentation on Climate Change and were given a list of 10 facts about this topic which they had to memorize and incorporate in their presentation. The specific topic was only revealed once they received the fact sheet.

The task could be divided in five phases: task instruction, fact sheet reading, mental preparation, presentation and recovery. EEG was recorded during the mental preparation and recovery phase, and eye tracking was recorded during the presentation phase.

After the task instruction, participants were given 2 minutes to study the fact sheet followed by 5 minutes to mentally prepare their presentation. The fact sheet was not available during the preparation time and participants were not allowed to talk or take notes. Following the preparation time, participants completed the PASA questionnaire to assess anticipatory stress and could click to place the video call to the jury panel to start their presentation. Visual cues were used to mimic those of video calling services like Skype or FaceTime to sell the illusion that the video call was live. During the presentation, the test leader scored the amount of facts from the fact sheet that were included in the presentation. In addition, if participants froze or stopped short of the 5 minute mark the test leader would provide the amount of time that was left and urged them to fill the time as best as they could. If participants ran out of material, the test leader could also give predefined content prompts (e.g. What have you done to impact or benefit the environment?). After 5 minutes of presentation time, the experiment continued automatically to a 5 minute recovery phase where participants had to sit in silence and fixate on a fixation cross. All stimuli were presented using Tobii Studio, on a Tobii T120 monitor.

### Materials

#### Baseline Self-Report

Measures of key traits underlying the current tasks (i.e. anxiety, stress, memory and imagery) were taken to assess a priori group differences, as well as a baseline measure for state anxiety. Specifically we assessed state and trait anxiety levels using the State Trait Anxiety Inventory (STAI-S and STAI-T; Spielberg et al., 1970), trait worry using the Penn State Worry Questionnaire (PSWQ; Meyer et al., 1990), stress reactivity using the Perceived Stress Scale (PSS; Cohen et al., 1983), and vividness of mental imagery using the Vividness of Visual Imagery Questionnaire (VVIQ; McKelvie, 1995). Scores for all questionnaires were computed using their respective scoring manual.

#### Task-Appraisal Self-Report

A second battery of questionnaires was administered right after the recovery phase of the TSST to assess post event state anxiety and task appraisals. For state anxiety we again used the STAI-S.

For post-event task appraisals, we used a combination of the visual analogue scales that are part of the Primary Appraisal and Secondary Appraisal scale (PASA) and items that were designed specifically for this study. The first part of the PASA was administered right before the presentation phase of the TSST, and consists of 16 items on anticipatory stress that are rated on a 6 point Likert-scale (1 strongly disagree – 6 strongly agree).

The second part of the PASA is administered after the stress-inducing event (in this case the TSST presentation) and consists of four visual analogue scales (0 not at all – 100 totally) on the level of experienced stress during and level of experienced control over the event. We elaborated on this scale with 7 more items. Of the novel items, two related to the level of physical and emotional discomfort (“The past situation was embarrassing for me” and “I felt physically uncomfortable in the past situation”) and two to confidence in their own performance (“I think my presentation went well” and “If I were asked to give the presentation again I would do it exactly the same way”). These items were rated on the same scale as the original PASA items (i.e., 0 not at all – 100 totally).

Furthermore, three items assessed subjective appraisal of the jury panel. Of these three, one used the original scale (“I thought the jury panel was intimidating”) and two were rated from negative to positive (0 – 100; “I think the judges evaluated my presentation” and “I felt the facial expressions of the jury panel were”). After reverse scoring positively worded items, all ratings were summed and averaged with the three jury-items forming their own category.

#### TSST video

In the original protocol for the TSST, the presentation participants give is held in front of a live audience of judges that are in the same room with the participant. However, several video based adaptations of this procedure have been developed over the years to accommodate the use of specific measures or manipulations. These adapted versions, event animated VR environments, are generally found to be equally effective as the original in-vivo setup. Here, we opted to use a pre-recorded video of the jury panel both to standardize the experience between participants and to accommodate the recording of EEG and eye-tracking during the TSST.

Three actors were recruited as jury members. Actors were instructed to wear professional attire. The recording started with a brief introductory statement *(“Welcome, thank you for preparing this presentation. You may start now*.*”*) by the head jury member, seated in the middle, to sell the idea of a live video connection. This was followed by 5 minutes of silent observing as if the jury were actually listening to a presentation. Following the official TSST protocol the jury was asked to refrain from giving verbal or non-verbal feedback during the entirety of the recording. However, taking notes and attentive listening were encouraged. Jury members looked directly into the camera to make it look like they were making eye contact from the participants perspective. After 5 minutes the head jury member notified the participants that their presentation time was up and that they would disconnect the call *(“Well time is up, thank you for your presentation. We will now close this connection*.*”*). Following this statement, one of the other jury members pressed a button on their laptop and the video switched to a black screen.

### Eye tracking recording and pre-processing

Eye tracking was recorded using a Tobii T120. Calibration was done right before the task instructions of the TSST, using a 9 point fixation calibration. Points were re-calibrated when the deviation was outside of the circle radius to get as clean of a measurement as possible. If deviations were still too large after three calibrations on the majority of the fixation points, or if the eye-tracker had trouble locating the pupil, testing was ceased. No chin or head rest was used as this would have interfered too much with the participants ability to speak during the presentation phase of the TSST. However, due to the cable setup of the EEG participants were highly restricted in their head movements.

Pre-processing of the eye tracking signal was performed in Tobii Studio. First, participants were rejected if gaze detection was less than 60 percent during the presentation phase. All further recordings were manually checked for quality by the lead experimenter. For the remaining dataset, areas of interest (AOI) were set as ellipses around the faces of the three jury members. The AOI’s were manually centered on each face throughout the recording to correct for movements of the jury members. A secondary group of AOI were set on the hands of the jury members, which were again adjusted if any movement occurred. The number of fixations and fixation duration (in ms) were calculated for all AOI’s and non-AOI (fixation on neither hands nor faces) using the automatic detection mechanism in Tobii Studio. The average fixation duration and total amount of fixations on each area were calculated per 1 minute interval to assess changes in fixation behavior throughout the 5 minute presentation.

### EEG recording and software

EEG recording was done using 64 Ag/AgCl electrodes placed in an extended 10-20 montage and collected at a 1,024 Hz sampling rate using the ActiveTwo BioSemi system (BioSemi, The Netherlands). Biosemi Common Mode Sense (CMS) active electrode and Driven Right Leg (DRL) passive electrode replaced the conventional ground electrode, and CMS was used as the online reference. Vertical EOG was measured with two Ag-AgCl electrodes placed above and below the right eye.

### EEG pre-processing and analyses

Offline pre-processing of the EEG time series was performed using MATLAB (The Mathworks, Version 9.6.0.1472908, R2019a) with EEGLAB (Version 2020). The EEG signal was down-sampled to 512 Hz. Data was re-referenced to the average of all 64 electrodes and offline band-pass filtered between 1 and 40 Hz (24dB/oct), with a 50-Hz notch filter (zero-phase shift). Noisy channels were interpolated. Ocular artefacts were removed from the non-resting state data (mental preparation and recovery) using the ocular correction ICA method in EEGLAB. For each condition, data was then segmented into 8-second non-overlapping epochs (4,096 time samples), to have sufficient low-frequency cycles to detect dPAC (Aru et al., 2015; Cohen, 2014). The first and last 15 epochs of both tasks were manually inspected for gross artifacts and excluded if necessary. Out of those, three early and three late clean epochs were selected for use in further analysis. These three early and three late epochs of the resting state, mental preparation, and recovery were exported to ASCII files for further analyses.

The focus of this study includes frontally mediated delta-beta PAC, to allow comparisons with relevant studies (Harrewijn et al., 2016; Poppelaars et al., 2018; Putman et al., 2012) and to reduce the risk of the multiple comparisons problem. F3, Fz and F4 were selected for further analysis, as these electrodes were the electrodes of interest of this study to allow comparisons with prior relevant studies on this topic (cf., Harrewijn et al., 2016; Poppelaars et al., 2018; Putman et al., 2012).

PAC analyses were performed as in Poppelaars et al. (2018) using custom scripts. The selected EEG epochs were down-sampled to 128 Hz, and band-pass filtered separately for delta (1–4 Hz) and beta (14–30 Hz) using a Butterworth IIR bandpass filter by using a zero phase-shift filtering method (with a filter order of 8 for delta and 34 for beta; which doubled after using both a forward and a backward filter). A Hilbert transform was applied to the delta and beta filtered epochs to isolate the phase and amplitude information (Papoulis & Pillai, 2000). The first and last 16 samples – equal to the order of the lower frequency’s filter (cf., Knyazev, 2011) – were cut from each epoch to remove edge artefacts originating from filtering (Aru et al., 2015; Kramer, Tort, & Kopell, 2008).

### Phase-amplitude coupling analysis

PAC analyses between delta phase and beta amplitude were performed using the debiased PAC (dPAC) method (Cox et al., 2014; Van Driel et al., 2015) with custom-written scripts (as in Poppelaars et al., 2018), to fit the current data specifications and research interests. Delta-beta dPAC and the accompanied Z-values were calculated for each participant and electrode, over the six epochs, and were thereafter averaged over the three electrodes, yielding one dPAC and Z-value per participant, per condition. dPAC was calculated by removing the phase clustering from the traditional PAC method (cf., Canolty et al., 2006) via a simple linear subtraction (cf., Cox et al., 2014; Van Driel et al., 2015). PAC can be defined as:

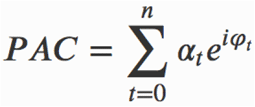

where a_t_ represents the amplitude of the modulated frequency (i.e., beta amplitude), and φ_t_ represents the phase of the modulating frequency (i.e., delta phase), *t* is time, and *n* is the total number of time samples. The phase clustering (PC) is calculated by averaging the complex vector of phase angles (*e*^*iφt*^), from which the magnitude (or strength) and angle of clustering can be determined:

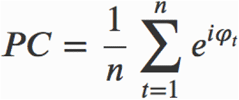

It should be noted that by not including the beta amplitude *a*, all complex numbers have the same length, and, therefore, all angles have the same weight in the averaging process. This allows for determining the average angle, or PC. For dPAC, the aforementioned complex numbers, *a*_*t*_*e*^*iφ*^ (combining beta amplitude *a* and delta phase *φ*) are averaged for all time samples, correcting the phase angle of the complex numbers by the earlier obtained PC:

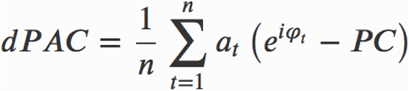

The dPAC value is expressed as the magnitude of the averaged complex number, where zero indicates no coupling, and values greater than zero indicate coupling. The significance of the coupling was established by comparing the dPAC values to surrogate dPAC values that were obtained via a non-parametric permutation testing approach (Maris & Oostenveld, 2007) by randomly shuffling epochs for phase information, while amplitude remained intact. This shuffling process was repeated 1,000 times, yielding a distribution of surrogate dPAC values as expected under the null hypothesis of no coupling. This method not only allows for significance testing but also accounts for possible outliers (Van Driel et al., 2015). Significant dPAC was determined by comparing dPAC to their surrogate counterparts (*dPAC*_*null*_) to obtain *Z*-values (dPACz):

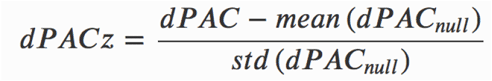

These *Z*-values were used for hypothesis testing due to their straightforward interpretation (i.e., standard deviation units; Cohen, 2014).

## Acknowledgements

The authors would like to thank Vanessa Danzer, Marie Dhoop and Mallissa Watts for their assistance with data collection, and Bart Endhoven, Martin van der Ploeg and Leanne van Est for acting as the jury members in the TSST video. This study was supported with a Vici innovational research grant from the Netherlands Organization for Scientific Research (NWO 453-15-005) awarded to IME.

## Author Contributions

NDM, LG and IME designed the study; NDM collected the data; NDM, DvS and LG analyzed the data; all authors interpreted the data; NDM wrote the first draft of the article, and DvS, LG and IME provided critical revisions. All authors approved the final version of the manuscript for submission.

## Ethics approval

The study was approved by the institutional ethical review board at Utrecht University (FETC19-053).

## Data and Code availability

Materials, code and data for all outcome measures generated during and/or analyzed during the current study will be made openly available upon publication.

## REFERENCES

Addis, D. R., Wong, A. T., & Schacter, D. L. (2008). Age-related changes in the episodic simulation of future events. Psychological Science, 19(1), 33–41. https://doi.org/10.1111/j.1467-9280.2008.02043.x

Angelidis, A., Hagenaars, M., van Son, D., van der Does, W., & Putman, P. (2018). Do not look away! Spontaneous frontal EEG theta/beta ratio as a marker for cognitive control over attention to mild and high threat. Biological Psychology, 135, 8–17. https://doi.org/10.1016/j.biopsycho.2018.03.002

Audrain, S., & McAndrews, M. P. (2020). Schemas provide a scaffold for neocortical integration at the cost of memory specificity over time. BioRxiv. https://doi.org/10.1101/2020.10.11.335166

Banks, S. J., Eddy, K. T., Angstadt, M., Nathan, P. J., & Phan, K. L. (2007). Amygdala-frontal connectivity during emotion regulation. Social Cognitive and Affective Neuroscience, 2(4), 303–312. https://doi.org/10.1093/scan/nsm029

Benoit, R. G., Szpunar, K. K., & Schacter, D. L. (2014). Ventromedial prefrontal cortex supports affective future simulation by integrating distributed knowledge. Proceedings of the National Academy of Sciences of the United States of America, 111(46), 16550–16555. https://doi.org/10.1073/pnas.1419274111

Bjärehed, J., Sarkohi, A., & Andersson, G. (2010). Less positive or more negative? Future-directed thinking in mild to moderate depression. Cognitive Behaviour Therapy, 39(1), 37–45. https://doi.org/10.1080/16506070902966926

Boland, J., Riggs, K. J., & Anderson, R. J. (2018). A brighter future: The effect of positive episodic simulation on future predictions in non-depressed, moderately dysphoric & highly dysphoric individuals. Behaviour Research and Therapy, 100, 7–16. https://doi.org/10.1016/j.brat.2017.10.010

Brooker, R. J., Phelps, R. A., Davidson, R. J., & Goldsmith, H. H. (2015). Context differences in delta beta coupling are associated with neuroendocrine reactivity in infants. Developmental Psychobiology, 58(3), 406–418. https://doi.org/10.1002/dev.21381

Buschman, T. J., & Miller, E. K. (2007). Top-down versus bottom-up control of attention in the prefrontal and posterior parietal cortices. Science, 315(5820), 1860–1862. https://doi.org/10.1126/science.1138071

Butler, G., & Mathews, A. (1987). Anticipatory anxiety and risk perception. Cognitive therapy and research.

Campbell, K. L., Madore, K. P., Benoit, R. G., Thakral, P. P., & Schacter, D. L. (2018). Increased hippocampus to ventromedial prefrontal connectivity during the construction of episodic future events. Hippocampus, 28(2), 76–80. https://doi.org/10.1002/hipo.22812

Campbell-Sills, L., Simmons, A. N., Lovero, K. L., Rochlin, A. A., Paulus, M. P., & Stein, M. B. (2011). Functioning of neural systems supporting emotion regulation in anxiety-prone individuals. Neuroimage, 54(1), 689–696. https://doi.org/10.1016/j.neuroimage.2010.07.041

Canolty, R. T., & Knight, R. T. (2010). The functional role of cross-frequency coupling. Trends in Cognitive Sciences, 14(11), 506–515. https://doi.org/10.1016/j.tics.2010.09.001

Chen, J., van den Bos, E., & Westenberg, P. M. (2020). A systematic review of visual avoidance of faces in socially anxious individuals: Influence of severity, type of social situation, and development. Journal of Anxiety Disorders, 70, 102193. https://doi.org/10.1016/j.janxdis.2020.102193

Cho, S., White, K. H., Yang, Y., & Soto, J. A. (2019). The role of trait anxiety in the selection of emotion regulation strategies and subsequent effectiveness. Personality and Individual Differences, 147, 326–331. https://doi.org/10.1016/j.paid.2019.04.035

Clark, D. A., & Beck, A. T. (2010). Cognitive theory and therapy of anxiety and depression: convergence with neurobiological findings. Trends in Cognitive Sciences, 14(9), 418–424. https://doi.org/10.1016/j.tics.2010.06.007

Cohen, S., Kamarck, T., & Mermelstein, R. (1983). A global measure of perceived stress. Journal of Health and Social Behavior, 24(4), 385–396. https://doi.org/10.2307/2136404

Demblon, J., Bahri, M. A., & D’Argembeau, A. (2015). Neural correlates of event clusters in past and future thoughts: How the brain integrates specific episodes with autobiographical knowledge. Neuroimage, 127, 257–266. https://doi.org/10.1016/j.neuroimage.2015.11.062

Devitt, A. L., & Schacter, D. L. (2018). An optimistic outlook creates a rosy past: the impact of episodic simulation on subsequent memory. Psychological Science, 29(6), 936–946. https://doi.org/10.1177/0956797617753936

Dijkstra, N., Mazor, M., Kok, P., & Fleming, S. (2021). Mistaking imagination for reality: Congruent mental imagery leads to more liberal perceptual detection. Cognition, 212, 104719. https://doi.org/10.1016/j.cognition.2021.104719

Ditta, A. S., & Storm, B. C. (2016). Thinking about the future can cause forgetting of the past. Quarterly Journal of Experimental Psychology, 69(2), 339–350. https://doi.org/10.1080/17470218.2015.1026362

Engel, A. K., & Fries, P. (2010). Beta-band oscillations--signalling the status quo? Current Opinion in Neurobiology, 20(2), 156–165. https://doi.org/10.1016/j.conb.2010.02.015

Frühholz, S., Godde, B., Lewicki, P., Herzmann, C., & Herrmann, M. (2011). Face recognition under ambiguous visual stimulation: fMRI correlates of “encoding styles”. Human Brain Mapping, 32(10), 1750–1761. https://doi.org/10.1002/hbm.21144

Gilboa, A., & Marlatte, H. (2017). Neurobiology of Schemas and Schema-Mediated Memory. Trends in Cognitive Sciences, 21(8), 618–631. https://doi.org/10.1016/j.tics.2017.04.013

Gilboa, A., & Moscovitch, M. (2017). Ventromedial prefrontal cortex generates pre-stimulus theta coherence desynchronization: A schema instantiation hypothesis. Cortex, 87, 16–30. https://doi.org/10.1016/j.cortex.2016.10.008

Harrewijn, A., Van der Molen, M. J. W., & Westenberg, P. M. (2016). Putative EEG measures of social anxiety: Comparing frontal alpha asymmetry and delta-beta cross-frequency correlation. Cognitive, Affective & Behavioral Neuroscience, 16(6), 1086–1098. https://doi.org/10.3758/s13415-016-0455-y

Herten, N., Otto, T., & Wolf, O. T. (2017). The role of eye fixation in memory enhancement under stress - An eye tracking study. Neurobiology of Learning and Memory, 140, 134–144. https://doi.org/10.1016/j.nlm.2017.02.016

Hutchinson, J. B., & Turk-Browne, N. B. (2012). Memory-guided attention: control from multiple memory systems. Trends in Cognitive Sciences, 16(12), 576–579. https://doi.org/10.1016/j.tics.2012.10.003

Jing, H. G., Madore, K. P., & Schacter, D. L. (2016). Worrying about the future: An episodic specificity induction impacts problem solving, reappraisal, and well-being. Journal of Experimental Psychology: General, 145(4), 402–418. https://doi.org/10.1037/xge0000142

Jing, H. G., Madore, K. P., & Schacter, D. L. (2017). Preparing for what might happen: An episodic specificity induction impacts the generation of alternative future events. Cognition, 169, 118–128. https://doi.org/10.1016/j.cognition.2017.08.010

Kirschbaum, C., Pirke, K. M., & Hellhammer, D. H. (1993). The “Trier Social Stress Test”: A tool for investigating psychobiological stress responses in a laboratory setting. Neuropsychobiology, 28(1-2), 76–81. https://doi.org/119004

Knyazev, G. G. (2007). Motivation, emotion, and their inhibitory control mirrored in brain oscillations. Neuroscience and Biobehavioral Reviews, 31(3), 377–395. https://doi.org/10.1016/j.neubiorev.2006.10.004

Knyazev, G. G. (2011). Cross-frequency coupling of brain oscillations: an impact of state anxiety. International Journal of Psychophysiology, 80(3), 236–245. https://doi.org/10.1016/j.ijpsycho.2011.03.013

Knyazev, G. G., Savostyanov, A. N., & Levin, E. A. (2005). Uncertainty, anxiety, and brain oscillations. Neuroscience Letters, 387(3), 121–125. https://doi.org/10.1016/j.neulet.2005.06.016

Kret, M. E., Denollet, J., Grèzes, J., & de Gelder, B. (2011). The role of negative affectivity and social inhibition in perceiving social threat: an fMRI study. Neuropsychologia, 49(5), 1187–1193. https://doi.org/10.1016/j.neuropsychologia.2011.02.007

Landkroon, E., van Dis, E. A. M., Meyerbröker, K., Salemink, E., Hagenaars, M. A., & Engelhard, I. M. (2021). Future-oriented Positive Mental Imagery Reduces Anxiety for Exposure to Public Speaking. Behavior Therapy. https://doi.org/10.1016/j.beth.2021.06.005

Liu, B., Wang, Y., & Li, X. (2018). Implicit emotion regulation deficits in trait anxiety: an ERP study. Frontiers in Human Neuroscience, 12, 382. https://doi.org/10.3389/fnhum.2018.00382

Masís-Obando, R., Norman, K. A., & Baldassano, C. (2021). Schema representations in distinct brain networks support narrative memory during encoding and retrieval. BioRxiv. https://doi.org/10.1101/2021.05.17.444363

Mather, M., & Sutherland, M. R. (2011). Arousal-Biased Competition in Perception and Memory. Perspectives on Psychological Science, 6(2), 114–133. https://doi.org/10.1177/1745691611400234

Mathews, A., & MacLeod, C. (2002). Induced processing biases have causal effects on anxiety. Cognition & emotion, 16(3), 331–354. https://doi.org/10.1080/02699930143000518

McKelvie, S. J. (1995). The VVIQ as a psychometric test of individual differences in visual imagery vividness: a critical quantitative review and plea for direction. Journal of Mental imagery.

Meyer, T. J., Miller, M. L., Metzger, R. L., & Borkovec, T. D. (1990). Development and validation of the Penn State Worry Questionnaire. Behaviour Research and Therapy, 28(6), 487–495. https://doi.org/10.1016/0005-7967(90)90135-6

Miloyan, B., Bulley, A., & Suddendorf, T. (2016). Episodic foresight and anxiety: Proximate and ultimate perspectives. The British Journal of Clinical Psychology, 55(1), 4–22. https://doi.org/10.1111/bjc.12080

Paulus, P. C., Dabas, A., Felber, A., & Benoit, R. G. (2021). Simulation-based learning influences real-life attitudes. https://doi.org/10.31234/osf.io/k8nxm

Pile, V., Williamson, G., Saunders, A., Holmes, E. A., & Lau, J. Y. F. (2021). Harnessing emotional mental imagery to reduce anxiety and depression in young people: an integrative review of progress and promise. The Lancet. Psychiatry, 8(9), 836–852. https://doi.org/10.1016/S2215-0366(21)00195-4

Poppelaars, E. S., Harrewijn, A., Westenberg, P. M., & van der Molen, M. J. W. (2018). Frontal delta-beta cross-frequency coupling in high and low social anxiety: An index of stress regulation? Cognitive, Affective & Behavioral Neuroscience, 18(4), 764–777. https://doi.org/10.3758/s13415-018-0603-7

Poppelaars, E. S., Klackl, J., Pletzer, B., & Jonas, E. (2021). Delta-beta cross-frequency coupling as an index of stress regulation during social-evaluative threat. Biological Psychology, 160, 108043. https://doi.org/10.1016/j.biopsycho.2021.108043

Putman, P. (2011). Resting state EEG delta-beta coherence in relation to anxiety, behavioral inhibition, and selective attentional processing of threatening stimuli. International Journal of Psychophysiology, 80(1), 63–68. https://doi.org/10.1016/j.ijpsycho.2011.01.011

Renner, F., Murphy, F. C., Ji, J. L., Manly, T., & Holmes, E. A. (2019). Mental imagery as a “motivational amplifier” to promote activities. Behaviour Research and Therapy, 114, 51–59. https://doi.org/10.1016/j.brat.2019.02.002

Ryan, J. D., & Shen, K. (2020). The eyes are a window into memory. Current Opinion in Behavioral Sciences, 32, 1–6. https://doi.org/10.1016/j.cobeha.2019.12.014

Schacter, D. L., Benoit, R. G., & Szpunar, K. K. (2017). Episodic future thinking: mechanisms and functions. Current Opinion in Behavioral Sciences, 17, 41–50. https://doi.org/10.1016/j.cobeha.2017.06.002

Schutter, D. J. L. G., & Knyazev, G. G. (2012). Cross-frequency coupling of brain oscillations in studying motivation and emotion. Motivation and emotion, 36(1), 46–54. https://doi.org/10.1007/s11031-011-9237-6

Schutter, D. J. L. G., Leitner, C., Kenemans, J. L., & van Honk, J. (2006). Electrophysiological correlates of cortico-subcortical interaction: a cross-frequency spectral EEG analysis. Clinical Neurophysiology, 117(2), 381–387. https://doi.org/10.1016/j.clinph.2005.09.021

Spielberg, C. D., Corsuch, R. L., & Lushene, R. E. (1970). The state-trait anxiety inventory (STAI). Test manual X form consulting psycologist.

Sylvester, C. M., Corbetta, M., Raichle, M. E., Rodebaugh, T. L., Schlaggar, B. L., Sheline, Y. I., Zorumski, C. F., & Lenze, E. J. (2012). Functional network dysfunction in anxiety and anxiety disorders. Trends in Neurosciences, 35(9), 527–535. https://doi.org/10.1016/j.tins.2012.04.012

Tse, D., Langston, R. F., Kakeyama, M., Bethus, I., Spooner, P. A., Wood, E. R., Witter, M. P., & Morris, R. G. M. (2007). Schemas and memory consolidation. Science, 316(5821), 76–82. https://doi.org/10.1126/science.1135935

van Kesteren, M. T. R., Rignanese, P., Gianferrara, P. G., Krabbendam, L., & Meeter, M. (2020). Congruency and reactivation aid memory integration through reinstatement of prior knowledge. Scientific Reports, 10(1), 4776. https://doi.org/10.1038/s41598-020-61737-1

van Son, D., Schalbroeck, R., & Angelidis, A. (2018). Acute effects of caffeine on threat-selective attention: moderation by anxiety and EEG theta/beta ratio. Biological ….

Wang, S.-H., & Morris, R. G. M. (2010). Hippocampal-neocortical interactions in memory formation, consolidation, and reconsolidation. Annual Review of Psychology, 61, 49–79, C1. https://doi.org/10.1146/annurev.psych.093008.100523

Williams, J. M., & Broadbent, K. (1986). Autobiographical memory in suicide attempters. Journal of Abnormal Psychology, 95(2), 144–149. https://doi.org/10.1037//0021-843x.95.2.144

